# Genetic screens identify connections between ribosome recycling and nonsense mediated decay

**DOI:** 10.1101/2021.08.03.454884

**Authors:** Karole N. D’Orazio, Laura N. Lessen, Anthony J. Veltri, Zachary Neiman, Miguel Pacheco, Raphael Loll-Krippleber, Grant W. Brown, Rachel Green

## Abstract

The decay of messenger RNA with a premature termination codon (PTC) by nonsense mediated decay (NMD) is an important regulatory pathway for eukaryotes and an essential pathway in mammals. NMD is typically triggered by the ribosome terminating at a stop codon that is aberrantly distant from the poly-A tail. Here, we use a fluorescence screen to identify factors involved in NMD in *S. cerevisiae*. In addition to the known NMD factors, including the entire UPF family (UPF1, UPF2 and UPF3), as well as *NMD4* and *EBS1*, we identify factors known to function in post-termination recycling and characterize their contribution to NMD. We then use a series of modified reporter constructs that block both elongating and scanning ribosomes downstream of stop codons and demonstrate that a deficiency in recycling of 80S ribosomes or 40S subunits stabilizes NMD substrates. These observations in *S. cerevisiae* expand on recently reported data in mammals indicating that the 60S recycling factor ABCE1 is important for NMD (1, 2) by showing that increased activities of both elongating and scanning ribosomes (80S or 40S) in the 3’UTR correlate with a loss of NMD.

**Author Summary:** In this work, we aim to understand the mechanism of targeting mRNAs for decay via the long-studied nonsense mediated decay (NMD) pathway. We demonstrate that efficient large and small subunit ribosome recycling are necessary components of NMD. We go on to provide evidence that either scanning or actively translating ribosomes in the 3’UTR disrupt the decay of NMD targets. Our work highlights the importance of the composition of the 3’UTR in NMD signaling and emphasizes the need for this region to indeed be untranslated for NMD to occur. Exon junction complexes (EJCs) in the 3’UTR are known to induce NMD, however, in the budding yeast system used here, the NMD targets are EJC-free. Therefore, our data support a model in which factors other than EJCs may accumulate in the 3’UTR and provide a signal for NMD.

## Introduction

Nonsense-mediated decay (NMD) is a quality control pathway that targets mRNAs for decay when ribosomes encounter an early or “premature” termination codon (PTC) (3). PTCs can arise from errors in the nucleus such as mis-splicing events or mutations during DNA replication and transcription, or from errors in translation such as use of alternative initiation sites or ribosome frameshifting (4). NMD also plays a broad regulatory role in eukaryotes by targeting both functional, alternatively spliced isoforms, and processed mRNAs that leave the nucleus but that do not encode functional gene products (e.g. long non-coding RNAs) (5). In each scenario, NMD is signaled through ribosome-dependent stop codon recognition and the mRNA is rapidly decayed.

NMD depends on the UPF and SMG-related proteins in all systems (6–8). These factors are critical for recognition of terminating/recycling ribosomes at PTCs and for triggering the recruitment of RNA decay machinery. Specifically, the NMD-central RNA helicase Upf1 interacts directly with eRF1 and eRF3 (9, 10) and this interaction is modulated by the phosphorylation status of Upf1 in mammalian systems (11, 12). At the same time, eRF3 interacts with the highly conserved NMD factors Upf2 and Upf3 to stabilize a complex of Upf1, Upf2, and Upf3 (13). What is likely essential to understanding the mechanism and specificity of NMD is an understanding of how the ribosome, and in turn release factors and recycling factors, distinguishes between normal and premature termination codons.

The normal processes of translation termination and recycling have been robustly characterized and begin when the 80S ribosome encounters a stop codon (UAA, UAG, or UGA) in the A site. The complex of eRF3:eRF1 recognizes the three stop codons and eRF3 then deposits eRF1 into the A site, in a manner analogous to eEF1A loading aminoacyl-tRNAs there during elongation (14). Following release of eRF3, the ATPase Rli1 (or ABCE1 in mammals) binds to the ribosome to promote termination (15) and ribosome recycling (15, 16) in which the 60S subunit dissociates from the complex composed of the 40S subunit, mRNA and P-site tRNA. Hcr1, a loosely bound member of the eIF3 initiation complex, has also been implicated in recruitment of Rli1 to termination sites and in 80S recycling (17, 18). In the final steps of recycling, the 40S subunit is dissociated from the tRNA and mRNA in a reaction promoted by three proteins known as Tma64, Tma20 and Tma22 in yeast (or eIF2D, MCT-1 and DENR in mammals) (19–22).

During any of the steps of termination and recycling, the activity of the ribosome at the stop codon can in principle signal NMD. What then are the features that distinguish between recognition of normal termination codons (NTCs) and premature termination codons (PTCs) that lead to NMD? In broad terms, the “context” of a stop codon within an mRNA determines whether the mRNA is targeted for NMD and could include: (1) different nucleotide sequences that affect recruitment of termination and recycling factors, (2) proximity to the poly-A tail, and (3) the composition and context of local RNA binding proteins. While much is known about how these different models might dictate NMD, we reasoned that important players in the pathway might remain undiscovered and could shed light on molecular mechanism.

Here, using a bi-directional fluorescent reporter in *S. cerevisiae*, we screen for factors that contribute specifically to decay of the mRNA or its translation repression during NMD. Along with the known NMD regulators in yeast, we identify a group of genes involved in translation termination and recycling. We find that deletion of the known 40S recycling factors *TMA20* and *TMA22* leads to the stabilization of NMD substrates and, similar to recent results describing the role of ABCE1 in NMD in mammals (1), we find that Hcr1 deletion leads to stabilization of NMD substrates. To test whether translocating ribosomes downstream of stop codons are responsible for disrupting NMD, we use reporter constructs containing different ‘barriers’ downstream of the PTC to block 80S translation and 80S/40S scanning and show that NMD-promoted decay of these reporters is restored in recycling deficient backgrounds. Taken together, these data support a model in which both actively translating 80S ribosomes as well as empty, scanning 80S ribosomes and 40S subunits disrupt critical signals for EJC independent NMD in the 3’ UTR.

## Results

### Developing a reporter system for nonsense mediated RNA decay

To identify genes necessary for NMD in yeast, we designed gene constructs that would report on both mRNA level and translation repression but not on nascent peptide stability. To implement this, we used a GFP-His3 conjugated protein and inserted a viral 2A peptide to effectively dissociate the GFP reporter protein from the downstream His3 protein that encodes a PTC. To control for overall expression, we used a bi-directional, inducible galactose promoter that expressed RFP in the opposite direction from GFP-2A-His3, and we normalized to RFP expression. This reporter system contains a FLAG epitope at the N terminus of His3 for detection of the peptide downstream of the 2A signal (Figure 1A). In the His3 ORF, we added a PTC to create an NMD signal (the NMD reporter) while the control reporter (the OPT reporter) contained no PTC (Figure 1A) (23). As anticipated for an NMD reporter construct, we find that insertion of a PTC ∼400 bp upstream from the normal stop codon of His3 leads to a 3-fold decrease in GFP/RFP levels as determined by flow cytometry and a 2-3-fold decrease in RNA as determined by northern blot analysis (Figure 1B and 1C).

**Figure 1:**
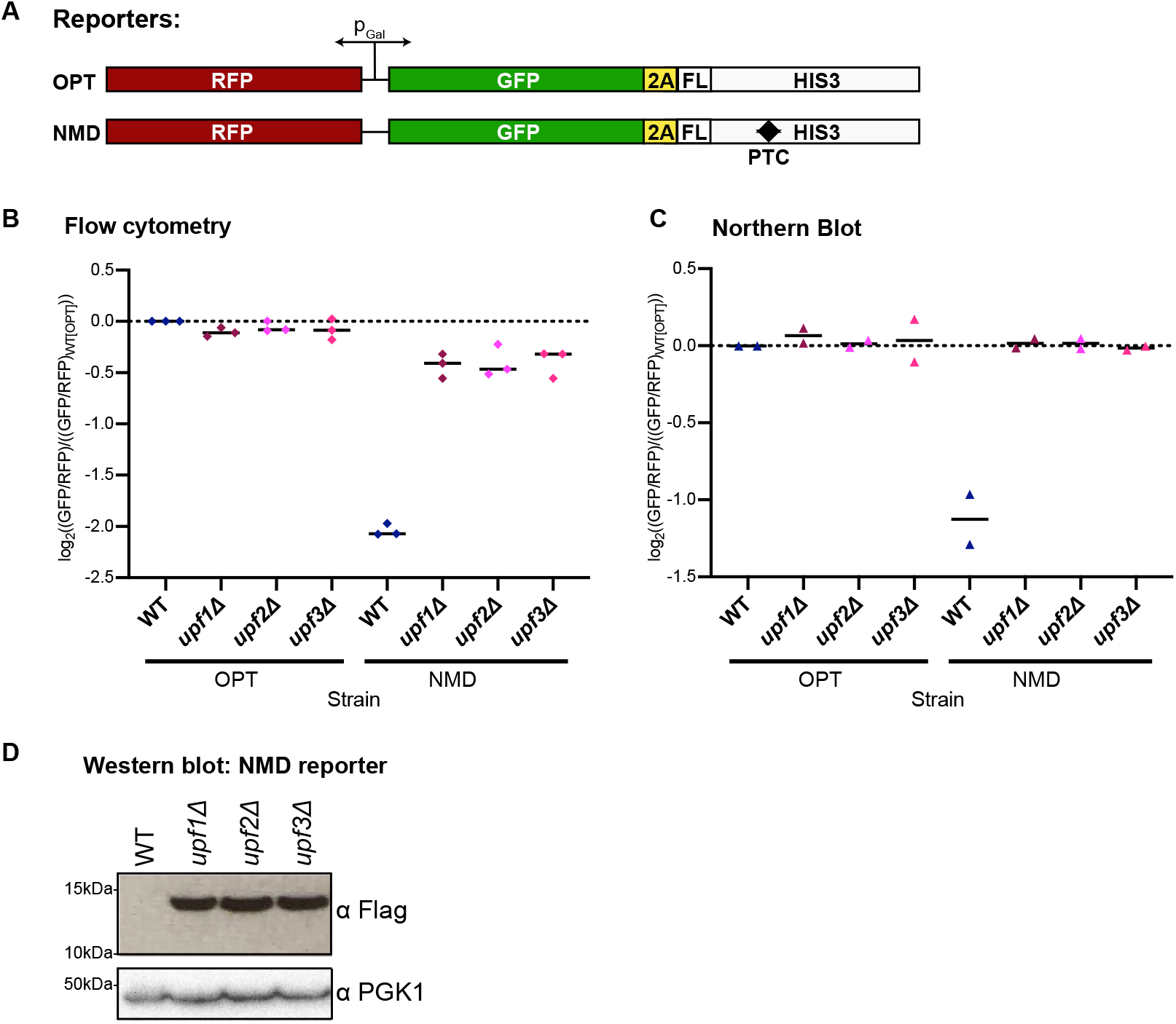
Fluorescent reporters reflect mRNA levels of an NMD target. (A) Schematic of reporters with a bi-directional, galactose-inducible promoter. RFP is in one direction, and GFP tethered to His3 via the 2A peptide and flag tag (FL) is in the other direction. (B) Individual averages of three different flow cytometry experiments performed on the indicated strains with the indicated reporters are shown. The GFP/RFP signal for each given strain normalized to the WT strain with the OPT reporter is plotted. (C) Northern blot quantification of two unique northern blots is shown. The GFP/RFP signal for each strain, normalized to the WT strain with the OPT reporter is plotted. (D) Western blot of the flag-tagged NMD reporter in the given backgrounds is shown with PGK1 as a control.

To confirm that the decreases in RNA levels for the reporter reflect NMD, we deleted the core NMD genes, *UPF1*, *UPF2* and *UPF3*, and saw a restoration of RNA levels for the NMD reporter by both flow cytometry and northern blots (Figure 1B and 1C). As expected, loss of these factors have no effect on the OPT reporter. Interestingly, in the NMD-reporter strains we saw that GFP RNA levels were fully restored by deletion of *UPF1/2/3* but GFP protein levels were not (compare Figure 1B and 1C), consistent with translation repression targeting NMD mRNAs (24), even in the absence of the UPF proteins. We next utilized the FLAG epitope to look at the peptide product of the NMD reporter. In WT cells, this FLAG epitope was undetectable via western blot as the mRNA is degraded by NMD and very little peptide is made. In the *UPF1/2/3* deletion strains this peptide construct was stabilized to a greater extent than the 3-fold stabilization seen for RNA (Figure 1D compared to 1B and 1C). These data are consistent with active degradation of the nascent peptide accompanying NMD, as reported previously (25).

### Screen to identify factors involved in NMD

We next performed a high-throughput reverse genetic screen using the *S. cerevisiae* reporter-synthetic genetic array (R-SGA) method (26, 27). In R-SGA, the reporter haploid strain is crossed with an array of viable haploid budding yeast deletion strains. Then, using a series of selections, an array of haploid strains containing the reporter and a single gene deletion is produced, allowing reporter activity to be scored in each deletion mutant. We performed independent screens with the OPT reporter (screen 1) and NMD reporter (screen 2) strains, where the reporter/deletion arrays were maintained on glucose media, then the cells were shifted to galactose to induce reporter expression. The GFP and RFP signal from the arrays were evaluated by fluorimetry, yielding a readout for both the OPT and NMD reporter (Suppl. Figure 1A and 1B) (23). For each strain in the plate array, Z-score normalized ratios of GFP intensity over RFP intensity were calculated. In this setting Z-scores represent the deviation of the GFP/RFP ratio for a given strain from the mean GFP/RFP ratio of the entire array; these data for the OPT and NMD reporter strains are plotted against one another in Figure 2A (raw data is given in Suppl. Table 1 and 2). From the NMD screen, there were 76 hits with a Z-score above 2.0 and 94 hits with a Z-score below −2.0 (Figure 2B, duplicate strains are removed in the Venn diagram). The OPT reporter screen, which has been previously published in D’Orazio et al. 2019, also included 64 hits with a Z-score above 2.0 and 138 hits with a Z-score below −2.0, with 27 genes overlapping as hits in both the OPT and NMD screens (Figure 2B) (23). However, GO enrichment data show that the hits in the NMD screen are strongly enriched in mRNA metabolic processing and nuclear-transcribed mRNA catabolic process nonsense-mediated decay, while there were no GO terms for the hits in the OPT screen with a similar level of enrichment (Figure 2B). Furthermore, *UPF1/2/3* deletion strains exhibited some of the largest deviations from the mean in the NMD reporter strains, immediately validating the potential of the screen (Figure 2A). Interestingly, *NMD4* and *EBS1* deletions are also strong outliers that cause an increase in GFP signal for the NMD reporter (Figure 2A). Nmd4 and Ebs1 are homologs of the Smg6 and Smg5/7 proteins in mammals which are key players in the nucleolytic decay of mammalian NMD substrates (28, 29) and the role of these factors in yeast has only recently been described (30).

**Figure 2:**
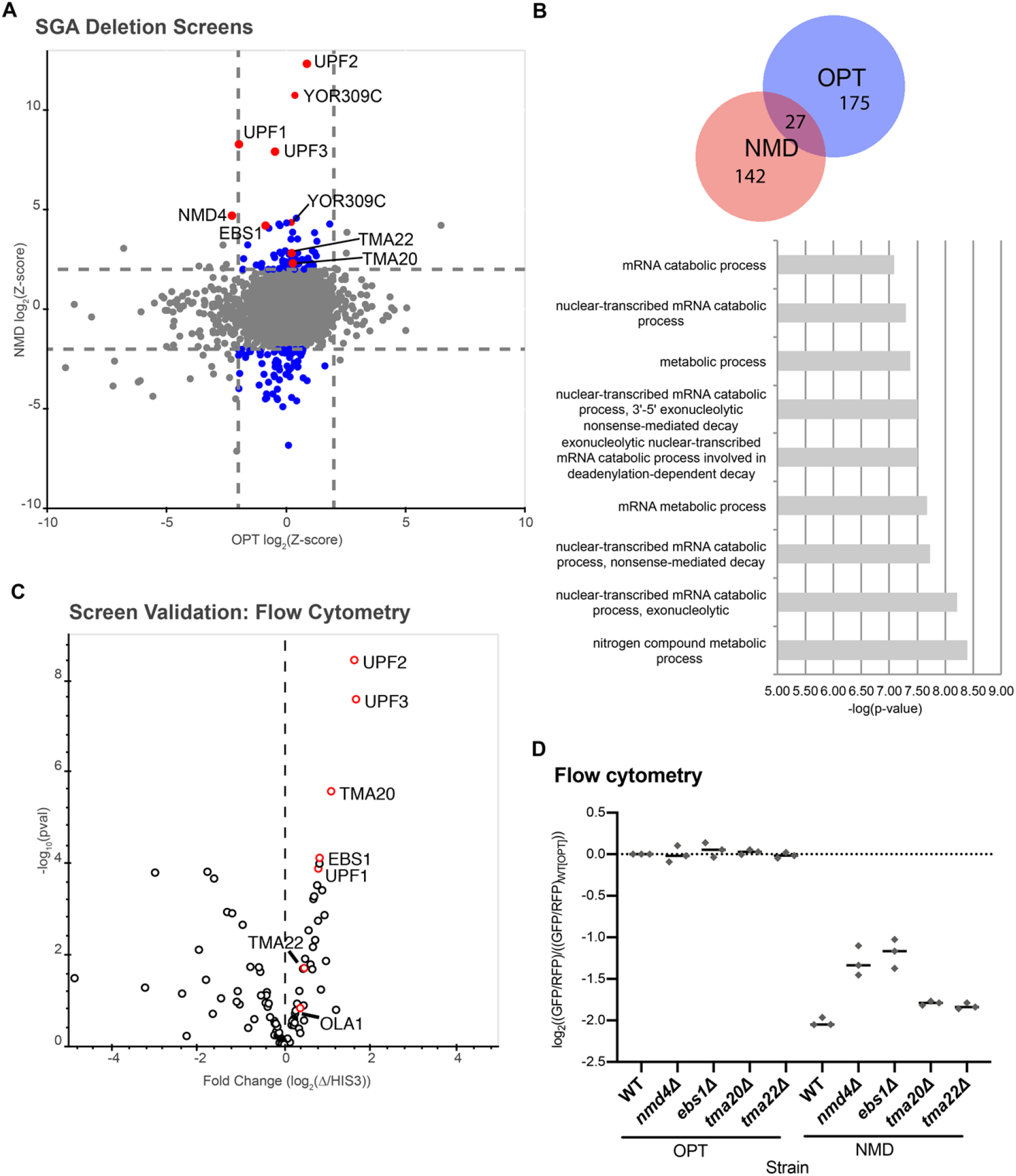
Yeast SGA screens identify genes important for RNA decay of NMD targets. (A) Plot of Z-scores from OPT reporter deletion screen versus the NMD reporter deletion screen. Z-scores reflecting the significance of log_2_(GFP/RFP) values from each deletion strain are plotted against each other for the two different GFP reporters. Dashed lines represent cutoffs at a Z-score greater than 2 or less than −2 for each reporter. Blue dots represent deletion strains that have a Z-score value outside the cut-off for the NMD reporter, but not for the OPT reporter. Red dots identify the strains known to affect NMD and/or strains of interest in this study. (B) (Top) Venn diagram showing the overlap between the OPT screen hits and the NMD screen hits. (Bottom) GO enrichment terms for cellular processes identified from the NMD hits performed using GOrilla software. Note: GO analysis for the OPT reporter hits did not give enrichment data. (C) Volcano plot showing data from a follow up screen using newly constructed yeast strains and flow cytometry. The x-axis compares the fold change of individual deletion strains to the control *HIS3* deletion strain. Data for each dot was obtained in triplicate and p-values are plotted on the y-axis. Red dots identify the strains known to affect NMD and/or strains of interest in this study. (D) Individual averages of three different flow cytometry experiments performed on the indicated strains with the indicated reporters are shown. The GFP/RFP signal for each given strain normalized to the WT strain with the OPT reporter is plotted.

As a validation, we re-transformed the NMD reporter into deletion strains from the deletion mutant collection corresponding to the NMD screen hits with a Z-score less than −2.0 or greater than 2.0. In this list, we excluded 29 genes that were also hits in the OPT screen, yielding 113 genes. We individually analyzed these 113 newly constructed strains by flow cytometry assays and plotted the fold change of each candidate gene deletion relative to a *HIS3*-deletion control (Figure 2C and Suppl. Table 3). Amongst the validated hits, along with the UPF proteins, Nmd4 and Ebs1, were two genes known to be involved in ribosome termination or recycling – *TMA20*, *TMA22* (20, 31).

There was one additional hit of interest, *YOR309C*, amongst the strongest hits of our NMD screen. Two individual strains with *YOR309C* deletions caused an increase in GFP (a number of deletion strains are duplicated in the SGA arrays). *YOR309C* is a “dubious open reading frame” that overlaps with the essential gene *NOP58* (Suppl. Figure 2A), which was shown to affect NMD in mammalian cells (32). As constructed, the *YOR309C* deletion results in a truncation of *NOP58* and uncharacterized effects on its 3’ UTR (due to insertion of a kanamycin cassette) (Suppl. Figure 2B). We therefore asked if truncation or loss of *NOP58* resulting from deletion of the *YOR309C* was responsible for blocking NMD that we observed in our screen (rather than from deletion of *YOR309C*). To test this, we complemented the *yor309cΔ (NOP58* truncated) strain with either full length Nop58 (Nop58-full) or the truncated version (Nop58-trunc) and evaluated the level of NMD by flow cytometry (Suppl. Figure 2C). Overexpression of either full length or truncated Nop58 restored NMD for the reporter (Suppl. Figure 2C). These data show that Nop58 truncation is not responsible for the NMD phenotype, consistent with studies demonstrating the functionality of Nop58 truncation proteins (33), and instead indicating that integration of KanMX at the *YOR309C* locus affects *NOP58* expression or Nop58 mRNA or protein stability.

We next decided to focus on genes that were thought to be involved in ribosome termination and/or recycling. We reconstructed the *TMA20-* and *TMA22-* deletions along with deletions for the previously known NMD-factors in the WT BY4741 strain carrying the OPT or NMD reporter and used these reconstructed strains for the rest of this study. We began by performing flow analysis on *nmd4Δ*, *ebs1Δ, tma20Δ, and tma22Δ* strains in triplicate. As seen recently, deletion of either *NMD4* or *EBS1* stabilizes the NMD reporter (Figure 2C) (30). Deletion of *TMA20* and *TMA22* also modestly stabilized the NMD reporter relative to the OPT reporter (Figure 2C), nicely underscoring the capacity of the R-SGA screens to isolate genes with even mild effects on NMD. Although the deletions showed mild effects on NMD, Tma20 and Tma22 have previously been shown to act redundantly, so we reasoned these mild effects might be indicative of a more significant combinatorial role in NMD for this group of proteins (19, 20).

### Efficient NMD requires efficient ribosome recycling

*In vitro* biochemical experiments and genetics combined with ribosome profiling experiments previously showed that Tma20 and Tma22 are involved in removal of the 40S subunit from mRNA following Rli1-mediated 60S dissociation (Skabkin *et al.*, 2010; Young *et al.*, 2018). These factors are homologous to the N- and C-termini of Tma64, which were shown to function redundantly with Tma20 and Tma22 in *in vitro* reconstituted systems that followed the release of 40S ribosomes and deacylated tRNA from mRNAs (20). Therefore, we asked if deletion of *TMA64* similarly causes stabilization of the NMD mRNA as observed in the *tma20Δ* and *tma22Δ* strains. Consistent with the screen data, where *TMA64* did not emerge as a candidate, and with recent data showing only a minor role for Tma64 *in vivo* (34), deletion of *TMA64* alone did not result in stabilization of the NMD reporter (Figure 3A). However, deletion of *TMA64* in combination with deletion of *TMA20* or *TMA22* did increase the GFP signal from the NMD reporter relative to each of the single deletions (Figure 3A). Northern blot data was broadly consistent with a general stabilization of the NMD reporter for the *TMA20*, *TMA22*, and *TMA64* deletion strains, although the double deletions (i.e. *tma20Δ tma64Δ*) did not show the enhanced effect as they did in the flow cytometry data (Figure 3B).

**Figure 3:**
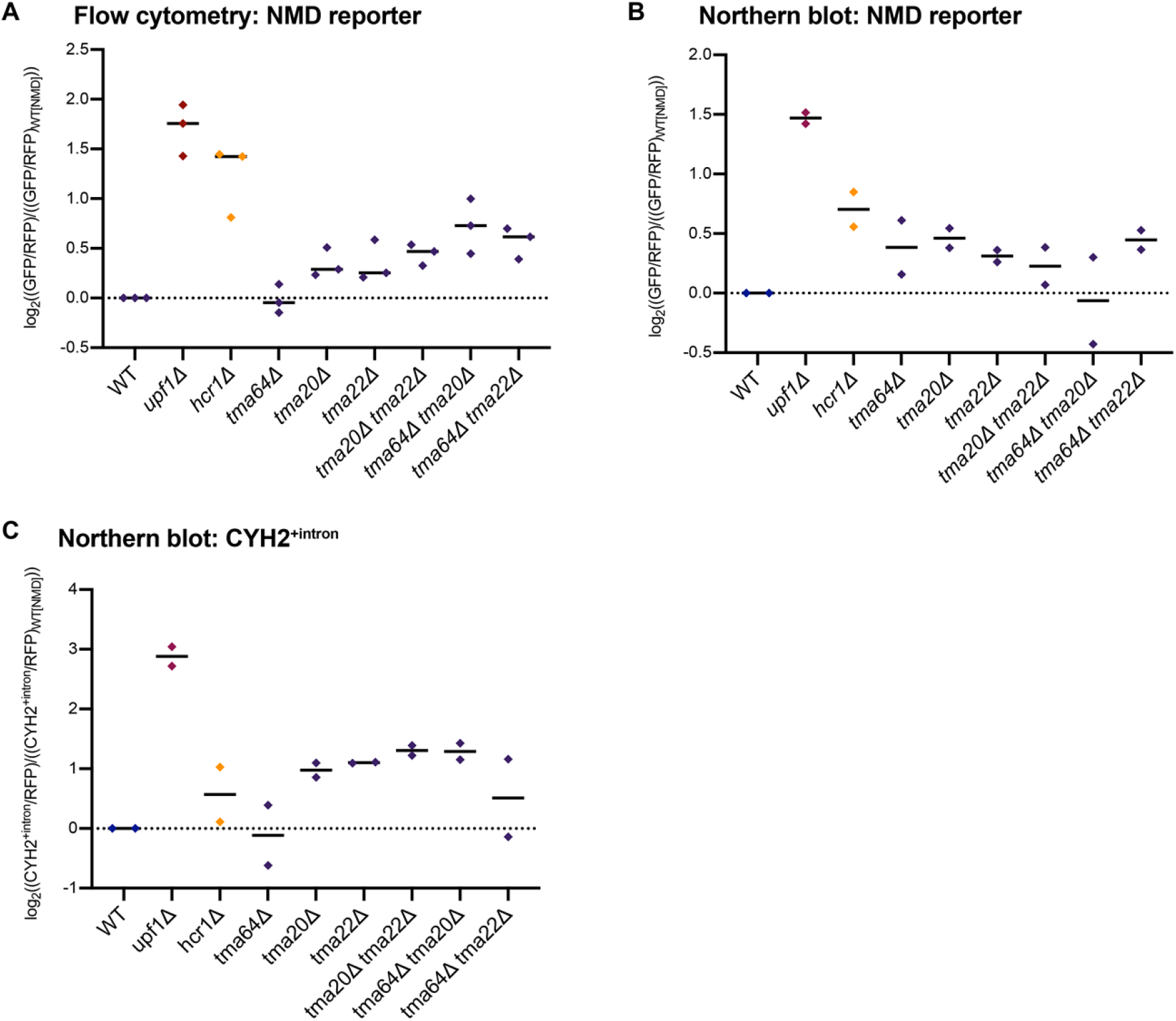
Deletion of both 40S and 60S ribosome recycling factors leads to stabilization of exogenous and endogenous NMD-targeted genes. (A) Individual averages of three different flow cytometry experiments performed on the indicated strains with the indicated reporters are shown. The GFP/RFP signal for each strain normalized to the WT strain with the OPT reporter is plotted. (B and C) Northern blot quantification of two unique northern blots is shown for each graph. The (GFP/RFP) signal in B and the (CYH2^+intron^/RFP) in C are both normalized to the WT strain with the OPT reporter and plotted.

Because *TMA20*, *TMA22*, and *TMA64* deletions have been shown to function in 40S ribosome recycling, we next asked if NMD was also affected by disrupting 60S ribosome recycling by deleting the Rli1-accessory factor gene *HCR1*. While the *hcr1Δ* strain did not survive the initial screen due to a general growth defect, this deletion indeed strongly stabilizes the NMD reporter (Figure 3A and 3B). These data together lead to the hypothesis that deficiencies in both 40S and 60S ribosome recycling can disrupt NMD.

We next assessed the effects of these same genes on a well characterized endogenous target of NMD, the rare intron-retaining transcript of *CYH2* (35). Deletion of the 40S recycling TMA factors stabilized this NMD substrate and deletion of *TMA64* enhanced the effect seen in both the *Δtma20* and *Δtma22* backgrounds, consistent with the reporter data (Figure 3C). Deletion of the 60S recycling factor *HCR1* had a significant, albeit weaker impact on NMD of the intron-containing *CYH2* transcript than that of the 40S recycling TMA factors. These data provide support for both the 40S and 60S recycling factors playing important roles in promoting efficient NMD. Taken together with previous studies, we reasoned that loss of these recycling factors could block NMD by allowing ribosomes to clear NMD-promoting factors from the 3’ UTR.

### Ribosomes in the 3’UTR disrupt NMD

As alluded to above, ribosome profiling experiments in Rli1 depletion and *hcr1Δ* strains or in *tma64Δtma20Δ* and *tma64Δtma22Δ* strains revealed an increase in the abundance of 80S ribosomes in 3’ UTRs transcriptome wide (19, 36). In the Rli1 depletion and *hcr1Δ* backgrounds, the failure of 80S termination or recycling increased ribosome readthrough and permitted an unusual 80S reinitiation process. These processes result in the accumulation of both elongating and scanning 80S ribosomes downstream of the stop codon (18, 36). In the *tma64Δtma20Δ and tma64Δtma22Δ* backgrounds, similar studies revealed an abundance of 80S ribosomes in the 3’ UTR, likely resulting from a lack of 40S recycling and subsequent reinitiation (19). Thus, failures in either recycling or termination could lead to the accumulation of ribosomes of various states in the 3’ UTR of mRNAs including: 1) translating 80S ribosomes; 2) non-translating 80S scanning ribosomes; or 3) 40S scanning ribosomes. In principle, these classes of ribosome are not mutually exclusive; for example, ribosomes that are actively translating in the 3’UTR can originate from scanning ribosomes that enter the 3’ UTR as a consequence of incomplete recycling and then reinitiate translation. To investigate the impact of such translocating ribosomes in the 3’ UTR on NMD, we designed a series of reporters with features that block the movement of ribosomes within the 3’ UTR subsequent to their encounter with the PTC and examined them in *hcr1Δ* and *tma64Δ tma20Δ* strains (Figure 4A).

**Figure 4:**
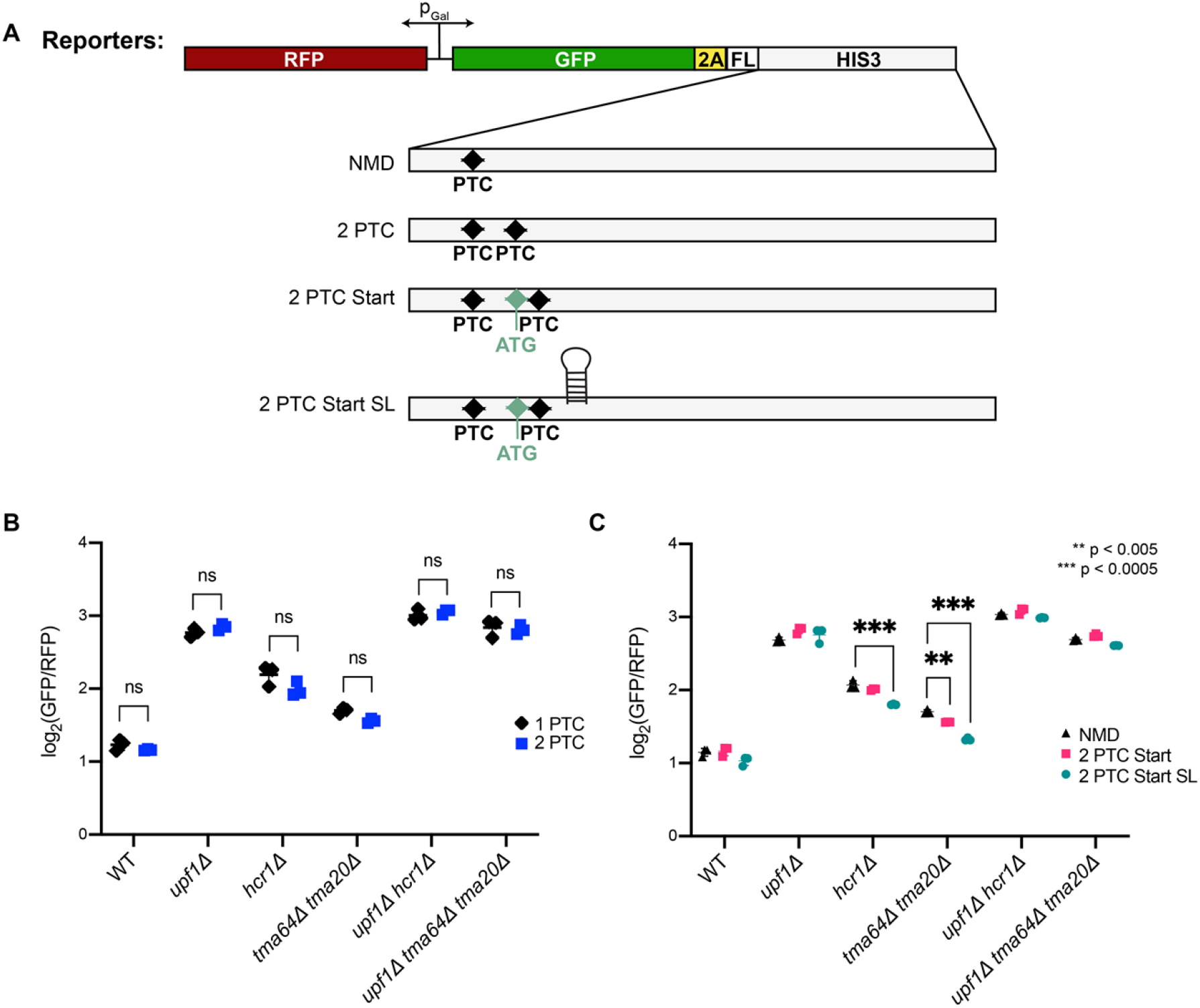
Ribosomes scanning or elongating in the 3’UTR impair NMD. (A) Schematic of reporters that alter specific characteristics of the 3’UTR of the original NMD reporter. Changes are made specifically to the region downstream of the premature stop codon. His3 is enlarged to highlight the additions. The ‘NMD’ reporter is the original NMD reporter with one PTC. ‘2 PTC’ has a second stop codon added downstream of the original PTC. ‘2 PTC Start’ includes a start codon immediately followed by a stop codon, both downstream of the original PTC. ‘2 PTC Start SL’ has a start codon immediately followed by a stop codon and then a stemloop directly downstream of this. (B and C) Individual averages of three different flow cytometry experiments performed on the indicated strains with the indicated reporters are shown. The GFP/RFP signal for each strain is plotted and significance is indicated with ‘ns’ = not significant, ‘**’ = p < 0.005, and ‘***’ = p < 0.0005.

The first reporter contained a second PTC <20 nucleotides downstream of the first PTC in the *HIS3* gene. If readthrough occurs at the first PTC, a statistically unlikely event, one would expect that upon encountering the next PTC, translating ribosomes would terminate protein synthesis and be recycled. If ribosome readthrough into the 3’ UTR is responsible for diminishing NMD in the various backgrounds, then NMD should be restored for this reporter (2 PTC). We observe that GFP levels in the *hcr1Δ* deletion strain and the *tma64Δ tma20Δ* strain are not affected by the addition of a second PTC (Figure 4B).

The absence of rescue of NMD that we observe with the ‘2 PTC’ reporter made us wonder whether scanning/non-translating 80S and/or 40S ribosomes might be instead be contributing to the diminished NMD that we observe in the various deletion strains. Accordingly, we asked if we could force scanning 80S to reinitiate through inclusion of a start-stop sequence in the 3’UTR that would then effectively clear ribosomes in the 3’ UTR by forcing them to terminate and recycle. The ‘Start + 2 PTCs’ reporter contains a start codon approximately 30 nt downstream of the PTC followed by a stop codon, with the remainder of the HIS3 sequence of the NMD reporter untouched (Figure 4A). In WT cells, addition of this sequence to the 3’UTR had no impact on reporter levels (Figure 4C). However, we also saw no effect of this sequence in the 3’UTR for the *hcr1Δ* strains and only a modest decrease in the *tma20Δ tma64Δ* strain, in which NMD was not completely restored (Figure 4C). These data suggested that either an unimpeded ribosome population could still be accessing the 3’ UTR and diminishing NMD, or that the ribosome recycling factors are affecting NMD through another mechanism.

In a final attempt to block ribosome access to the 3’ UTR, we added a highly structured stem loop structure to the 3’UTR of the ‘Start + 2 PTCs’ reporter to make the ‘Start + 2 PTCs + SL’ reporter. The selected stem loop sequence has been previously shown to block 40S scanning in the 5’ UTR and initiation (37); we recapitulate these earlier observations and show that when inserted into the 5’UTR of our reporter system, this stem loop sequence completely disrupts translation (Suppl. Figure 3A and 3B). Importantly, when inserted into the open reading frame of the OPT reporter, reporter levels are unchanged (Suppl. Figure 3A and 3B) consistent with ready unwinding of the stem loop by an elongating 80S. In WT cells, adding this stem loop sequence to the 3’UTR of our reporters has no impact on reporter expression (Figure 4C). However, in the *hcr1Δ* and *tma20Δ tma64Δ* backgrounds that are deficient in 60S and 40S recycling, respectively, the inserted stem loop sequence partially restores NMD (see decreased GFP/RFP ratios) (Figure 4C). Taken together, these data support a model (Figure 5) in which defects in ribosome recycling result in the accumulation of scanning ribosomes in the 3’UTR (both 40S and 80S), which effectively disrupts NMD.

**Figure 5:**
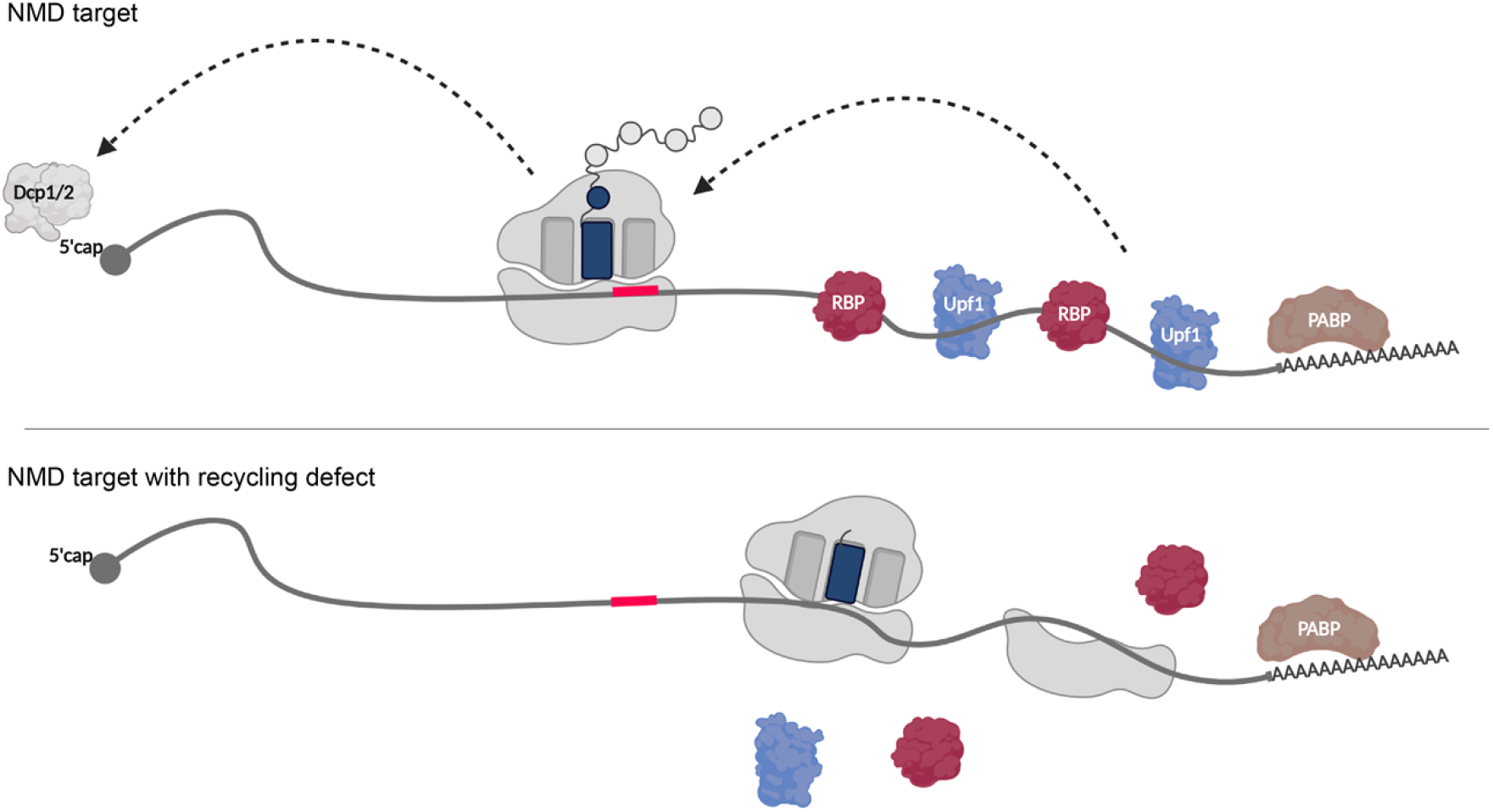
Model of how ribosomes acting in the 3’UTR affect NMD. Schematics of two NMD substrates are shown. On top, an mRNA in which normal ribosome recycling occurs is depicted with 3’UTR binding proteins downstream of the premature stop codon (red line). These factors are essential for recognition of a ribosome at an early stop codon and signal nonsense mediated decay through the decapping factors Dcp1/2. On the bottom, ribosome recycling is impaired causing ribosomes to scan through the stop codon, disrupt proteins binding in the 3’UTR, and disrupt the NMD pathway, which results in a stabilization of the mRNA. Created with Biorender.

## Discussion

Translation termination and recycling are inherently linked to NMD signaling through recognition of a premature stop codon by the ribosome. Many previous studies have worked to define how these ribosome activities, termination and recycling, might be connected to the specification of NMD either directly or through interaction with a host of factors potentially bound to the 3’ UTR of the mRNA (9,10,13,38–41).

Here we used a newly developed genetic screen in yeast with fluorescent reporters to identify key factors that contribute to NMD. We identify 40S ribosome recycling factors Tma20 and Tma22 in the screen, and additionally 60S recycling factor Hcr1 in follow up experiments, as important modulators of NMD both on reporter constructs and endogenous NMD targets (Figure 2 and 3). We then use a series of reporter constructs to establish that these ribosome recycling factors contribute to NMD by blocking ribosome activity in the 3’ UTR (Figure 4). These observations are consistent with early studies showing that ribosome reinitiation after a premature termination codon abrogated NMD (42, 43) and several recent studies in mammals implicating ribosome recycling and 3’ UTR ribosome activity in NMD specification (1, 2). Furthermore, these data nicely support models implicating 3’ UTR bound proteins as key determinants of NMD (38–41,44).

Rli1, or ABCE1 in mammals, works with its partner Hcr1 to stimulate termination and recycle 60S subunits (15,16,18,36). Our data show that deletion of Hcr1 stabilizes NMD substrates, consistent with recent work done in mammalian cells demonstrating that a loss of ABCE1 stabilizes NMD substrates (1, 2). However, the known dual role of Rli1/Hcr1 in both termination and recycling (15) confounds the interpretation of these results. We also found that deletion of 40S ribosome recycling factors – Tma64, Tma20, and Tma22 – causes a similar stabilization of NMD substrates (Figure 3). These data together suggest that any losses in ribosome recycling that lead to increased ribosome occupancy in the 3’ UTR could diminish NMD.

This model was tested using a series of reporters containing molecular perturbations that would block the movement of scanning or translating ribosomes. With an NMD reporter construct containing both a highly structured stem loop to block scanning ribosomes and stop codons to terminate re-initiated ribosomes in the 3’UTR, NMD was nearly completely restored in a 40S recycling-deficient background (*tma20Δ tma64Δ*) and partially restored in a 60S-recycling deficient background (*hcr1Δ*) (Figure 4). These data argue that efficient ribosome recycling is required to maintain a ribosome-free 3’UTR for robust NMD and suggest that 40S subunits in the 3’UTR can affect NMD.

Many previous studies have established that the 3’UTR is a critical regulator of NMD. In early studies in yeast, specific “downstream elements” (DSEs) were demonstrated to be important for signaling NMD and these DSE’s were later shown to be bound by the RNA-binding protein Hrp1 that was critical for NMD for a subset of DSE-containing targets (39, 44). In mammalian cells, the presence of EJCs and the accumulation of Upf1 in the 3’UTR have both been strongly implicated in signaling NMD (38,40,41). Taken together, these studies lead to simple models invoking positive contributions to NMD by proximal bound proteins (Upf1 and the EJC) that recruit other critical machinery involved in mRNA decay (such as the SMG proteins). The contributions of such proteins in the 3’UTR to NMD signaling provide an explanation for our results showing that the presence of scanning and/or translating ribosomes in the 3’UTR disrupts NMD, likely by displacing bound NMD promoting factors.

It is critical to reconcile these 3’ UTR-centric NMD models relative to other NMD models arguing that the actual efficiency of translation termination or recycling is critical for specifying NMD. In 2004, Amrani et al. showed that ribosome recycling is relatively inefficient at PTC’s compared to NTC’s and that tethering of a nearby PABP could rescue these recycling defects (45). Multiple studies have shown that PABP is a strong stimulator of efficient termination and subsequently argued that efficient termination is a negative determinant for NMD (46–50). What is currently missing is a causative link between the timing (kinetics) of termination or recycling and the induction of NMD.

Our results establish that deletion of factors that result in a loss of ribosome recycling decreases the likelihood of NMD. Moreover, the modifications that we made to the 3’ UTR reporter sequences to block ribosome movement increased NMD efficiency. Importantly, these modifications were unlikely to have impacted the efficiency of termination or recycling at the PTC. These constructs thus effectively function to separate ribosome activity at the stop codon from ribosome function post-termination/recycling. We interpret these data to mean that ribosomes moving in the 3’ UTR can displace proteins that are critical for NMD and that blocking their movement can rescue NMD (Figure 5). As a corollary, efficient termination/recycling catalyzed at NTC’s that are typically located proximal to the poly(A) tail may evade NMD not because termination/recycling is efficient but because there are few proteins in the short 3’ UTR to recruit the NMD decay machinery. Our data establish that efficient recycling is critical for NMD. However, the timing of termination/recycling could also be critical, so long as it proceeds to completion, in allowing for the recruitment of NMD promoting factors such as the UPF and SMG proteins. These ideas could potentially reconcile much of the seemingly contradictory literature in this area.

3’ UTRs are critical determinants of eukaryotic biology and 3’ UTR RNA binding proteins have been widely shown to be critical for RNA, localization, stability and translational control during development and beyond (51). Given that NMD is specified by recognition of aberrantly placed stop codons within an mRNA, the idea that the protein constituents bound to the nearby 3’ UTR contribute directly to the process makes considerable sense. Our results in the yeast system where there are no EJC’s, a major contributor to 3’UTR regulation of NMD in mammals, substantially broaden the NMD model that invokes contributions of 3’ UTR binding proteins (38,39,44).

## Materials and Methods

### Plasmid Construction

The OPT reporter plasmid, or pKD065, was constructed as described in D’Orazio et al 2019 (23). The NMD reporter plasmid (pKD081) was constructed by inserting an in frame ‘UAA’ stop codon 270 bp into the *HIS3* coding region of pKD065.

To make the ‘2 PTC’ and ‘2 PTC Start’ reporters used in Figure 4 in which pKD081 contains a second stop codon and pKD081 contains a start codon followed by a stop codon in all three reading-frames, respectively, pKD081 was first digested with XhoI and SpeI. Separately, pKD081 was PCR’d with primer sets KD001/KD002 and KD003/KD004 or primer sets KD001/KD011 and KD012/KD004. For each plasmid, the two PCR fragments and the digested plasmid were purified and assembled using Gibson Assembly. To make the ‘Scanning’ reporter used in Suppl. Figure 3 in which pKD065 contains a stem loop 10 bp upstream of the start codon, pKD065 was digested with AgeI and XhoI. Then pKD065 was PCR’d with primer sets KD005/KD006 and KD007/KD008. The two PCR fragments and the digested plasmid were then purified and assembled using Gibson Assembly. To make the ‘Elongation’ reporter used in Suppl. Figure 3 in which pKD065 contains a stem loop in the coding region of HIS3, pKD065 was digested with XhoI and SpeI. Then pKD065 was PCR’d with primer sets KD001/KD009 and KD010/KD004. The two PCR fragments and the digested plasmid were then purified and assembled using Gibson Assembly. Finally, to make the ‘2 PTC Start SL’ reporter used in Figure 4, pKD081 was digested with XhoI and SpeI. Then the ‘2 PTC Start’ reporter was PCR’d with KD001/KD013 and KD014/KD004. The two PCR fragments and the digested plasmid were then purified and assembled using Gibson Assembly. To make the Nop58 expressing plasmids, pRS316 was digested and full length and truncated *NOP58* including 500+ bp of the endogenous promoter were PCR’d. The PCR fragment was then inserted into the digested plasmid using Gibson Assembly.

### Yeast Strains and Growth conditions

Yeast strains used in this study are described in Suppl Table 4 and are derivatives of BY4741 unless specified otherwise. Yeast strains were constructed using standard lithium acetate transformations. Reporter strains were constructed by digesting the given plasmids with StuI and integrating the reporters into the *ADE2* locus of BY4741. Deletion strains were constructed by replacing the gene of interest with drug resistance cassettes at the given locus; see genotypes in Suppl Table 4.

For SGA experiments, query strains for the deletion screens were constructed by introducing the RFP---GFP-2A-FLAG-HIS3 cassettes from StuI digested pKD065, or pKD081 at the *ADE2* locus in Y7092 (26, 52) (Suppl. Table 4). For gal-induced growths, overnight cultures were grown in YPAGR media (Suppl. Table 4). Overnights were diluted in the same media to an OD of 0.1 and harvested at an OD of 0.4–0.5.

### Flow cytometry

Flow cytometry of individual strains was performed as in D’Orazio et al 2019. Briefly, cells were harvested in log phase and washed with PBS once, then ran on a Millipore guava easyCyte flow cytometer for GFP and RFP detection using 488 nm and 532 nm excitation lasers, respectively. Data for 10,000 cells were collected and gated based on size. Flow cytometry was done in triplicate, with each group of cells taken from individual growths. For triplicate plots, the average of each individual flow cytometry sample was taken and normalized to the indicated strain. The log of the fraction was then plotted for each experiment. For statistical analysis of flow cytometry triplicates, a standard t-test was run in prism and p-values are reported.

### Northern blots

Northern blots were performed as in D’Orazio et al 2019. Briefly, 25 ml of log-phase cells were harvested. RNA was isolated and 5 μg of RNA were loaded into a 1.2% agarose, formaldehyde denaturing gel and run for 1.5 hours. The RNA was vacuum transferred to a nitrocellulose membrane (N + Hbond, Amersham). The membrane was UV crosslinked, placed in pre-hybridization buffer and rotated at 42°C for an hour. The indicated DNA-oligo listed in Suppl. Table 4 was end labeled using gamma-ATP and T4 Polynucleotide Kinase radioactive labeling protocol from NEB. Labelled oligos were purified using GE Healthcare illustra ProbeQuant G-50 Micro Columns and the membrane was probed overnight, rotating at 42°C. The membrane was washed 3 times in 2 x SSC, 0.1% SDS for 20 min at 30°C, then exposed to a phosphoscreen. The phosphoscreen was scanned using a Typhoon FLA 9500.

### Western blots

Protein Isolation and western blotting were performed as discussed in D’Orazio et al 2019.

### Reporter SGA Screens

#### SGA procedure

SGA screens were performed using a Biomatrix Robot with a few modifications (S&P Robotics Inc.). Briefly, yKD176 and yKD179 query strains (Suppl. Table 4) were crossed individually with the yeast nonessential deletion library (26). The deletion library was arrayed in a 1536-format with 4 colonies per deletion strain. Because we found that our query strains had a slightly lower mating efficiency and growth defect, incubation times for every step of the SGA protocol were prolonged by 50-75%. Mating and sporulation steps were performed on standard SGA media (26, 52).

For each deletion screen, diploid strains were selected on DIP media then haploid double mutant strains were selected for multiple rounds on HAP media listed in Suppl. Table 4. Finally, to induce reporter expression, cells were pinned onto haploid double mutant selection medium with raffinose and galactose at a final concentration of 2% (HAPGR media listed in Suppl. Table 4). Cells were grown for 26-30 hours before scanning on a Typhoon FLA9500 (GE Healthcare) fluorescence scanner equipped with 488 nm and 532 nm excitation lasers and 520/40 and 610/30 emission filters. Plates were also photographed using a robotic system developed by S&P Robotics Inc. in order to determine colony size.

#### Reporter Screen Analysis

Screen analysis was performed as described in D’Orazio et al 2019 and previous manuscripts (23,53,54). Briefly, GFP and RFP fluorescent intensity data was collected using TIGR Spotfinder microarray software (Saeed AI, Sharov V, White J, Li J, Liang W, et al. TM4: a free, open-source system for microarray data management and analysis. Biotechniques. 2003;34:374–378). Colony size data was collected using SGATools (55) (http://sgatools.ccbr.utoronto.ca/). After border strains and size outliers (<1500 or >6000 pixels) were eliminated, median GFP and RFP values were taken. We then calculated log_2_(mean GFP/mean RFP) for the replicates and performed a LOESS normalization for each plate. Based on this LOESS normalized value, Z-scores were calculated. Strains for the NMD reporter with a Z-score greater than 2.0 or less than −2.0 were considered a hit if their Z-score in the OPT reporter was not also an outlier.

#### Venn Diagrams

The Venn diagram was created using BioVenn and the input was deletion strains with a Z-score greater than 2.0 or less than −2.0.

#### GO term analysis

Go analysis was performed with the data from the screen using a list of hits that were in the cutoff of Z > 2.0 or Z < −2.0 and the input gene list is the genes that GFP/RFP data was successfully acquired for in the deletion array screen. The P value cutoff was set to < 10^-7 using GOrilla (56, 57). Duplicate strains were removed during analysis.

#### Validation Screen

We selected NMD reporter genes that gave a Z-score greater than 2.0 or less than −2.0. Then we removed any hits that also had a Z-score greater than 2.0 or less than −2.0 for the OPT reporter.

These deletion strains were then struck out from a glycerol stock collection contributed by Brendan Cormack and transformed with the NMD reporter plasmids (pKD081). Three individual colonies from each transformation were isolated. Strains that did not grow were dropped from the experiment, yielding 101 deletion strains to test (Suppl. Table 3). These biological replicates were grown overnight in a flow cytometer plate in YPAGR and put on the flow cytometer the next day. The triplicate flow cytometry data for each strain was analyzed, normalizing to the HIS3 controls for all plates tested.

## Acknowledgments

We thank Brendan Cormack for his contributions to planning and carrying out experiments.

